# Quantifying cerebral blood arrival times using hypoxia-mediated arterial BOLD contrast

**DOI:** 10.1101/2022.03.27.485933

**Authors:** Alex A. Bhogal, Ece Su Sayin, Julien Poublanc, Jim Duffin, Joseph A. Fisher, Olivia Sobcyzk, David J. Mikulis

## Abstract

Cerebral blood arrival and tissue transit times are sensitive measures of the efficiency of tissue perfusion and can provide clinically meaningful information on collateral blood flow status. We exploit the arterial blood oxygen level dependent (BOLD) signal contrast established by precisely modulating arterial hemoglobin saturation using hypoxic respiratory challenges (dOHb-BOLD) to quantify arterial blood arrival times throughout the brain. A combination of hemodynamic lag with a modified carpet plot analysis yielded lag, onset (blood arrival), mean transit time (MTT) and hypoxic response information, which is indicative of relative total blood volume. Onset times averaged across 12 healthy subjects were 1.1 ± 0.4 and 1.9 ± 0.6 for cortical gray and deep white matter, respectively. The average whole brain MTT was 4.5 ± 0.9 seconds. The dOHb-BOLD response was 1.7 fold higher in grey versus white; in line with known differences in regional blood volume fraction. Our method was also applied in unilateral carotid artery occlusion patient, which revealed prolonged signal onset with normal perfusion in the affected hemisphere. In cases with exhausted reserve capacity or confounding flow effects such as vascular steal, dOHb-BOLD can potentially inform on collateral flow pathways to provide a valuable compliment to clinical vascular reactivity measures.

## 1.0 INTRODUCTION

Cerebral blood arrival and tissue mean transit times (BAT and MTT) show spatial variation in the healthy human brain. They are sensitive measures of the efficiency of tissue perfusion, potentially having the ability to assess the status and effectiveness of downstream vascular flow control. Vascular diseases alter BAT and TT such that mapping alterations of these metrics could prove useful in assessing the impact of vascular disease on the cerebral circulation in a clinically meaningful manner. More specifically, techniques that are sensitive to arterial hemodynamic status are of high clinical interest for their potential to identify the presence and effectiveness of collateral blood supply^1^. For instance, collateralization of leptomeningeal vessels is assumed to be a predictor of favorable outcome after stroke due to the ability to maintain perfusion in the ischemic penumbra for longer periods of time^2^. Similarly, an intact circle-of-Willis and/or strong leptomeningeal collaterals have been shown to protect against perfusion deficit in cases of significant carotid artery stenosis^3,4^. Cerebral auto-regulatory mechanisms play an essential role for capitalizing on alternate blood delivery pathways via vasodilatory responses that serve to draw blood towards affected tissues. Establishing the association between arterial collateralization and auto-regulatory capacity can provide a means to guide treatment decisions^5^ and potentially predict outcomes.

There are several Magnetic Resonance Imaging (MRI) based techniques available that can provide insights into arterial hemodynamics including arterial spin labeling (ASL), and dynamic susceptibility contrast (DSC) MRI. ASL is appealing in that it can provide quantification of cerebral blood flow (CBF). When combined with vasoactive stimuli such as hypercapnic breathing challenges or the injection of acetazolamide, pre- and post-stimulus data can be combined to reveal cerebrovascular reactivity (CVR). When acquired with a multi post labeling delay (multi-PLD) scheme^6^, ASL can hint at collateralization via the presence of arterial transit artifacts^7^ (ATA), prolonged arterial transit times (AAT^8^ or bolus arrival time (BAT^9^). However, ASL is a generally low signal-to-noise (SNR) modality with limited sensitivity in white matter (WM), and the temporal hemodynamic parameters rely on modeling the ASL kinetics^10^. DSC is also able to provide temporal information by tracking the passage of an injected contrast bolus using rapidly acquired gradient echo images. Using tracer kinetic modeling, perfusion measures as well as MTT and time to peak (TTP) can be derived, which may also reflect collateralization^11^ or impaired hemodynamics. Nonetheless. DSC-MRI may be considered invasive and relies on kinetic modelling implementation choices such as how to determine the arterial input function can lead to errors in hemodynamic parametrization.

CVR mapping based on Blood oxygen level dependent (BOLD-CVR) in combination with hypercapnic stimuli, has been proposed as a marker for recruitable collateral blood flow^12^. BOLD-CVR measures reflect local changes in the pattern of vascular resistance^13^ which is probed by the BOLD response using vasoactive stimulus. Such data however does not reflect the adequacy of baseline perfusion compensation. An additional caveat is that vasodilation can lead to changes in blood velocity that may modulate bolus arrival times^14^, or vascular steal and/or redistribution of blood flow that may confound true blood arrival and transit times in both healthy and disease affected tissues. In contrast to BOLD response to hypercapnia and in common with DSC, deoxy-hemoglobin can act as an arterial contrast agent^15,16^. By manipulating blood oxygen saturation at the lungs, hypoxia-BOLD (dOHb-BOLD) signal changes are reflective of both arteries and veins but without confounding effects related to altered hemodynamics.

To test this concept, 12 healthy subjects were scanned using a continuous BOLD acquisition throughout a transient hypoxia breathing paradigm. A combination of temporal delay mapping and a modified carpet plot analysis^17^ were applied to generate relative onset maps representing the arrival of the deoxy-hemoglobin bolus throughout the brain. In a single subject, comparison is made using a hypercapnic stimulus to reveal striking differences in the hemodynamic response. Finally, this method is tested in a patient with unilateral carotid artery occlusion to reveal extended onset times consistent with presumed collateral blood flow via the circle-of-Willis and/or leptomeningeal vessel network. We hypothesize that quantifying the onset of immediate signal changes induced by the arrival of a deoxy-hemoglobin bolus in brain tissues, provides a measure of arterial blood arrival time. Delays in this onset time can be taken to reflect longer or slower transit paths through the cerebral vasculature.

## 2.0 METHODS

### 2.1 Data acquisition

Data was selected from ongoing studies that are approved by the Research Ethics Board of the University Health Network and conform to the Declaration of Helsinki. Written informed consent was obtained from all subjects. MRI datasets from twelve healthy control subjects (33 ± 14 years old, 4 female) were analyzed. An additional patient with a unilateral carotid artery occlusion (CAO) was also included.

The study was performed using a 3-Tesla scanner (HDx Signa platform, GE healthcare, Milwaukee, WI, USA) with an 8-channel head coil. T2*-weighted gradient echo-planar imaging sequence with the following parameters: TE/TR = 30/1500 ms, flip angle = 73^⍰^, 29 slices, voxel size = 3 mm isotropic voxels and matrix size = 64 × 64. In the two cases in which a hypercapnic stimulus was applied (outlined below) a slightly modified BOLD acquisition was performed for increased spatial coverage: TR = 2400 ms, slices = 42, voxel size = 3.5 mm isotropic). A high-resolution T1-weighted spoiled-gradient-echo sequence was acquired for registration and tissue segmentation purposes with the following parameters: TI = 450 ms, TR 7.88 ms, TE = 3ms, flip angle = 12°, voxel size = 0.859 × 0.859 × 1 mm, matrix size = 256 × 256, 146 slices, field of view = 24 × 24 cm, no inter-slice gap

A gas blender with a sequential gas delivery breathing circuit (RespirAct, Thornhill Research, Canada) was used to deliver a double hypoxic bolus during isocapnea throughout the continuous BOLD acquisition. The hypoxic breathing protocol consisted of a 60 second baseline period in which end-tidal O_2_ (P_ET_O_2_) was targeted at 95 mmHg, followed by a 60 second step down to 40 mmHg, then a return to baseline for 20s targeting 95 mmHg, then a second step down again targeting 40 mmHg, and then a final return to baseline at 95 mmHg P_ET_O_2_. A measured P_ET_O_2_ trace is shown in figure 1B. In a single control, an additional CO_2_ challenge was applied consisting of clamping end-tidal partial pressure of CO_2_ (P_ET_CO_2_) at the subjects resting for 2 minutes, followed by a step increase by 10 mmHg for 2 minutes and returned to baseline for another 2 minutes. P_ET_CO_2_ was then reduced by 10mmHg for 1 minute, followed by a steady rise in P_ET_CO_2_ to 15mmHg above baseline over ~5 min, and returned to baseline for 2 minutes.

**Figure 1;.**
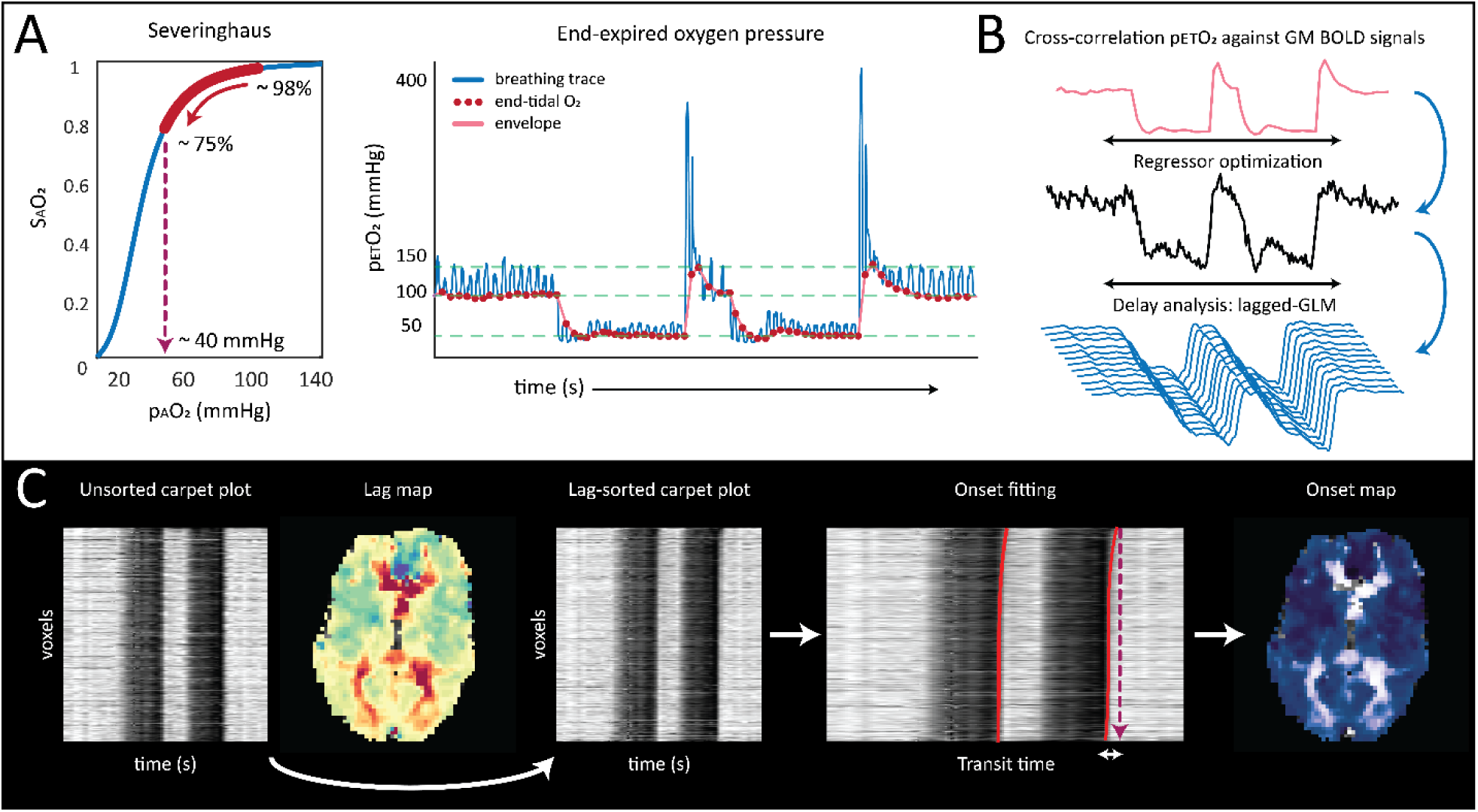
Process diagram for generating onset maps based on hypoxic breathing paradigm. (A): arterial oxygen tension is manipulated using a RespirAct system. Arterial oxygen pressure is targeted at 45 mmHg which, according to the Severinghaus equation^26^, leads to a predicted arterial hemoglobin saturation of approximately 75%; (B): resulting BOLD data, along with the end-tidal O_2_ trace are provided as input for the hemodynamic lag calculation algorithm. This process involves the generation of an optimized BOLD signal probe which is regressed against each voxel time-series at progressive lag increments; (C): the resulting hemodynamic lag values are then used to sort BOLD voxels in order on increasing lag. By plotting voxels as a list, 2D SHAG carpet plots are produced. Signal transitions driven by the BOLD signal onset are fit using a non-linear spline function and the resulting onset times are re-mapped into the native BOLD space to produce onset maps.

### 2.1 Removal of large vessel contributions

A thresholding step was applied to generate a modified whole-brain anatomical mask aimed at removing signal contributions from large veins that might overshadow arterial responses or strongly influence subsequent analysis steps. First, the inverse temporal signal to noise ratio (tNSR) was calculated for each voxel in the source BOLD data. Finally, voxels with tNSR values greater than the 98^th^ percentile were identified and voxels were removed from the input brain mask (see figure S1).

### 2.2 Lag mapping

Voxel-wise dOHb-BOLD data were linearly-detrended and temporally de-noised using a wavelet based approach^18^. Time-series were decomposed using the ‘sym4’ mother wavelet and signals were then de-noised by applying a hard Bayesian threshold to the resulting wavelet coefficients. Data were then spatially smoothed using a 3D Gaussian kernel (filter width: 5 voxels, FWHM: 5 mm). Processed dOHb-BOLD data and end-tidal O_2_ (P_ET_O_2_) data were interpolated by a factor 5 (effective TR: 300ms) to identify sub-TR signal displacements. Data were subsequently used to generate hemodynamic lag maps using a modified Rapidtide^19,20^ approach implemented in the Matlab-based seeVR toolbox^21^. In brief, a manual alignment between the end-tidal O_2_ (P_ET_O_2_) and average grey matter (GM) signal was performed to minimize alignment errors that may occur to noise and motion-related signal spikes when using automated methods. Next, P_ET_O_2_ traces were correlated with GM voxels (excluding large vessels) and voxels having a Pearson correlation higher than 0.7 were isolated and temporally aligned for principal component analysis. Principal components explaining at least 85% of the signal variability were extracted to generate a new regressor. The new regressor was then correlated against all GM voxels and this process was repeated until the root mean square error between subsequent regressor was less than 0.005. This convergence process resulted in an optimized dOHb-BOLD regressor. Hemodynamic lags were calculated using a shifted general linear model (GLM) approach that minimized the co-efficient of determination (R-squared) to select the shifted regressor that explained most of the dOHb-BOLD signal variance. Finally, a linear regression between the P_ET_O_2_ trace and each voxels’ signal trace was performed. The slope of this regression provided the hypoxic signal response.

### 2.3 Carpet plot analysis

Lag values were used to sort individual voxel time-series, which were then listed to produce 2-D carpet plots (figure 1C). The original work by Fitzgerald et. al^17^ applied the carpet plot analysis to derive the presumed transit times of blood borne signals observed in resting-state fMRI under the assumption of a linear relationship amongst voxels exhibiting progressive signal delays. In this work, we have expanded this model to account for non-linear onset behavior throughout the brain by characterizing signal ‘edges’ using a non-linear 3^rd^ order spline fit^21^. Moreover, we extended the carpet plot algorithm to map values back to the native functional space in order to generate spatial maps of blood arrival. Our reported mean transit times were derived using the original code provided by Fitzgerald et. al^17^. Briefly, the transit time is defined by the difference in onset time between the voxel with the longest lag (top of the lag-sorted carpet plot) and the voxel with the shortest lag (bottom of the lag-sorted carpet plot).

Based on our experience, the transition from hypoxia to normoxia occurred more rapidly that when going from normoxia to hypoxia. Blood re-oxygenation by the lung is faster than peripheral deoxygenation that is affected by slow transfer of dissolved oxygen from blood into the tissue. As a result, we chose to fit the onset edges around the two hypoxic bolus time-periods defined by the return to normoxia (see fig 1). Onset maps were calculated by averaging the voxel-wise onset values generated from both edges. We also averaged the two resulting transit time values for each subject. The same hemodynamic lag and carpet plot analysis was performed on data derived using hypercapnic step + ramp stimuli obtained in one of the healthy controls and also a patient with unilateral carotid artery stenosis. For the hypercapnic cases, the P_ET_CO_2_ was used as the input for the analysis described above. Due to the nature of the hypercapnic stimulus (i.e. a block followed by a ramp), the return to baseline P_ET_CO_2_ period was chosen for onset edge detection, since the progressive signal increase of the ramp stimulus did not provide sharp enough signal transitions.

### 2.4 Spatial normalization and statistical analysis

The mean BOLD image was spatially registered to the anatomical T1 scan using FSL’s^22^ FLIRT^23^ function and resulting transformation matrices were inverted. The anatomical image was brain extracted (BET^24^) and then segmented (FAST^25^) and the resulting tissue masks were transformed to BOLD space using the inverted transformation matrices; these masks were used in the analysis steps described in 2.1-2.2. The T1 image was then registered to the 2mm MNI atlas using linear (FLIRT) and non-linear (FNIRT^22^) registration. Transformation matrices were then applied to the hemodynamic parameters to normalize the individual lag maps, hypoxic response, BOLD-CVR maps, and onset maps to MNI space. MNI-registered data was averaged and mean and standard deviation of regional hemodynamic values were calculated using a series of MNI regions of interest (ROI) masks. Where reported, statistical significance (p<0.01) was investigated without the assumption of equal variance using a Welch’s *t*-test.

## 3.0 RESULTS

### 3.1 Group comparison

All subjects were able to complete the hypoxic protocol with no discomfort. Individual subject P_ET_O_2_ traces aligned with the corresponding average GM dOHb-BOLD signal response data are provided in figure S2A/B. Based on the Severinghaus equation^26^ (see also the equations of Balaban et al., 2013^27^), the arterial saturation changed from approximately 98% to 75% between arterial partial of oxygen (PaO_2_) of 95 mmHg to 40 mmHg (figure 1A). In general, the carpet plot analysis based on the hypoxic challenge delivered consistent onset/blood arrival maps across all 12 healthy subjects with values ranging between 0 and 4 seconds (figure 2). Compared with the MNI-averaged hemodynamic lag maps, MNI-averaged onset maps demonstrated less spurious values; for example, in deeper WM where blood volume and the related BOLD contrast to noise ratio (CNR) can be low (see figure 3).

**Figure 2;.**
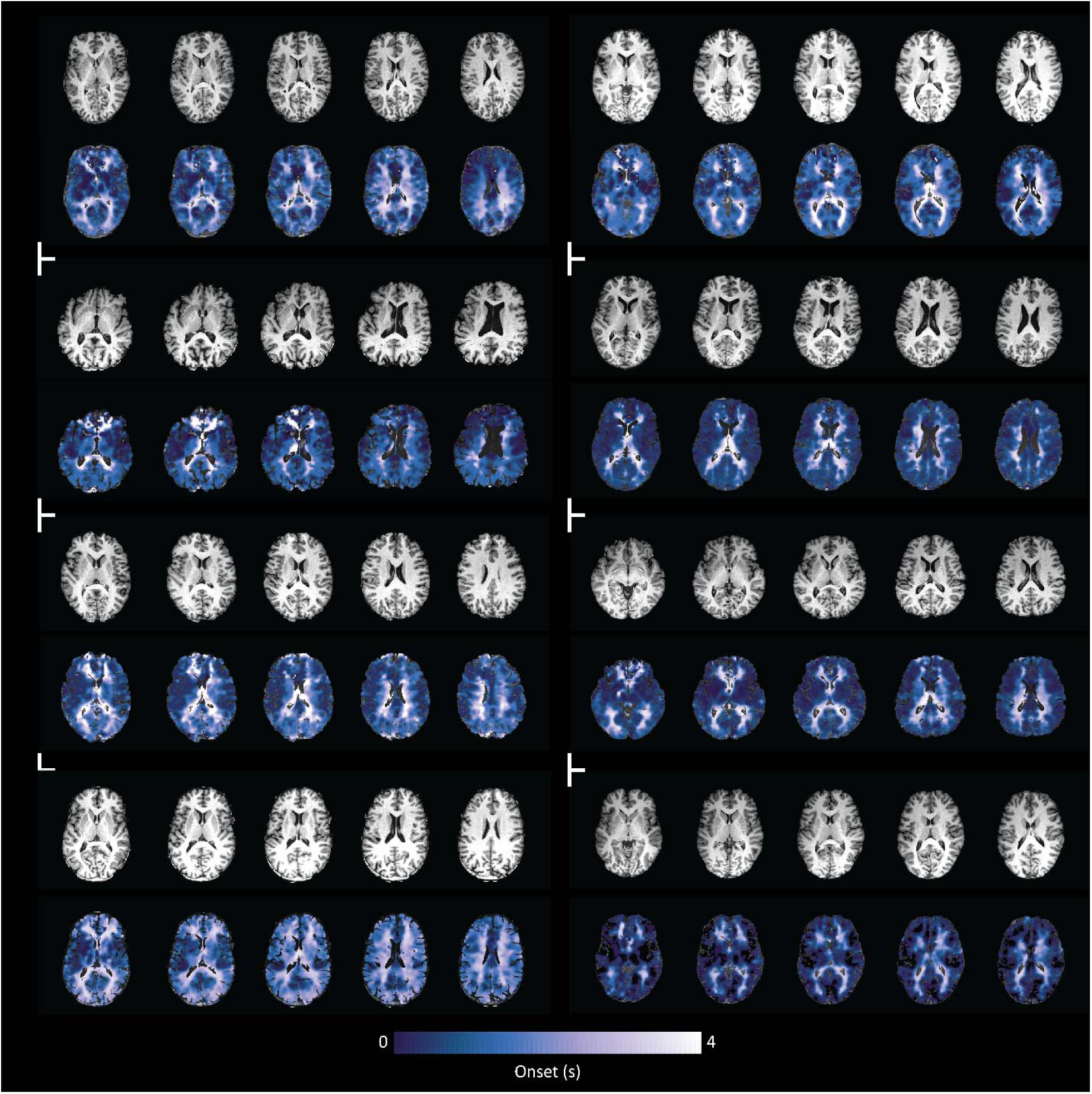
Representative onset maps normalized to anatomical T1 image for 8 subjects: hypoxic signal onset times are scaled between 0 and 4 seconds. Onset maps represent arterial arrival time and show remarkable inter-subject stability within this range. Associated P_ET_O_2_ traces and aligned GM responses for all subjects are provided in figure S2A and carpet plots are provided in figure S3A.

**Figure 3;.**
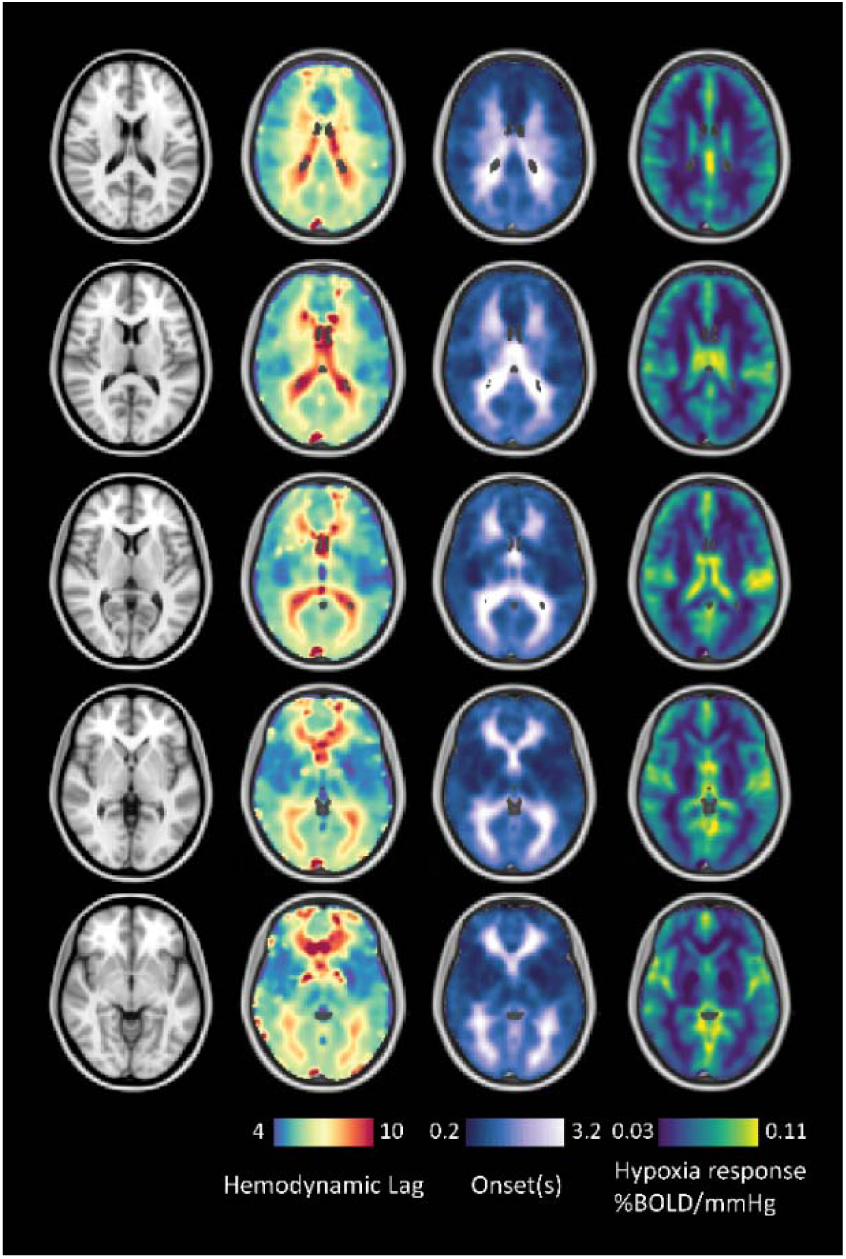
MNI averaged hemodynamic lag, onset and hypoxia signal response for 12 subjects: regional average parameter responses are provided in table 1. Average maps have been smoothed using a 3D Gaussian kernel with 4mm FWHM.

Mean and standard deviation lag, onset and hypoxic response values calculated in a series of MNI-ROIs are provided in table 1. Consistent with known differences in cerebral blood volume^28^, the hypoxic signal response in cortical GM was approximately 1.7-fold higher than that measured in the deep WM (0.05 ± 0.02 versus 0.03 ± 0.01 %ΔBOLD/mmHg P_ET_O_2_, respectively). Considering the assumption that the short hypoxic stimulus applied has limited effect on vascular tone or tissue metabolism, this distribution suggests that the regional magnitude of the hypoxic response may provide an indication of relative *total* blood volume (figure 3: right). When looking at regional onset times, the average onset of the occipital lobe was significantly longer than that of the frontal lobe (1.5 ± 0.4 versus 0.9 ± 0.4 %ΔBOLD/mmHg P_ET_O_2_, p < 0.001), which is in line with expectations considering that the posterior circulation may be delayed due to the smaller diameter of vertebral artery and other anatomic variations. In the original work of Fitzgerald et al.^17^, the mean transit time as calculated using the carpet plot technique, is assumed to represent the propagation time of signal patterns throughout the majority of voxels in the brain. The mean transit time (as calculated by averaging the values from both edges in our data: see supplemental figure 2) was 4.5 ± 0.9 seconds across twelve subjects.

**Table 1:** Mean and standard deviation calculated in MNI-averaged regions of interest (see figure 3). In the left column, MNI regions are sorted based on descending average lag values. On the right, MNI regions are sorted based on descending onset values. The hypoxic signal response is sorted in accordance with the onset times.

| Region (Lag-sorted) | Lag | Region (Onset-sorted) | Onset | H. Response |
| --- | --- | --- | --- | --- |
| caudate | $8.2 \pm 1.5$ | caudate | $2.0 \pm 0.6$ | $0.05 \pm 0.01$ |
| deep white matter | $7.8 \pm 0.7$ | deep white matter | $1.9 \pm 0.6$ | $0.03 \pm 0.01$ |
| white matter | $7.0 \pm 1.3$ | thalamus | $1.7 \pm 0.9$ | $0.07 \pm 0.02$ |
| cerebellum | $6.9 \pm 1.1$ | white matter | $1.6 \pm 0.8$ | $0.04 \pm 0.02$ |
| frontal lobe | $6.8 \pm 2.0$ | cerebellum | $1.5 \pm 0.3$ | $0.06 \pm 0.02$ |
| occipital lobe | $6.8 \pm 1.0$ | occipital lobe | $1.5 \pm 0.5$ | $0.05 \pm 0.01$ |
| thalamus | $6.8 \pm 1.3$ | hippocampus | $1.3 \pm 0.4$ | $0.06 \pm 0.02$ |
| temporal | $6.7 \pm 1.9$ | parietal lobe | $1.1 \pm 0.6$ | $0.05 \pm 0.01$ |
| cortical grey matter | $6.6 \pm 1.5$ | temporal | $1.1 \pm 0.5$ | $0.05 \pm 0.02$ |
| parietal lobe | $6.5 \pm 0.9$ | cortical grey matter | $1.1 \pm 0.4$ | $0.05 \pm 0.02$ |
| hippocampus | $6.4 \pm 0.9$ | insula | $0.9 \pm 0.5$ | $0.05 \pm 0.02$ |
| putamen | $5.8 \pm 0.8$ | frontal lobe | $0.9 \pm 0.4$ | $0.05 \pm 0.02$ |
| insula | $5.7 \pm 0.8$ | putamen | $0.9 \pm 0.4$ | $0.04 \pm 0.01$ |

### 3.2 Hypoxia versus hypercapnia

The average lag and onset values between GM and WM for the single subjects in which both protocols were performed were as follows: For hypercapnia, lag values were 15.8 ± 17.2 and 32.4 ± 23.4 seconds for GM and deep WM, respectively. Onset values were 4.0 ± 4.2 and 11.6 ± 9.0 seconds GM and WM, respectively. For hypoxia, lag values were 5.9 ± 3.9 and 6.0 ± 2.7 seconds for GM and deep WM, respectively. Onset values were 1.3 ± 0.8 and 1.5 ± 1.0 seconds for GM and deep WM, respectively. Differences in spatial coverage and acquisition parameters between hypercapnic and hypoxic acquisitions limit in-depth comparison. However the tendency towards longer response times for hypercapnia was visibly predominant (see figure 4A, S3B).

**Figure 4;.**
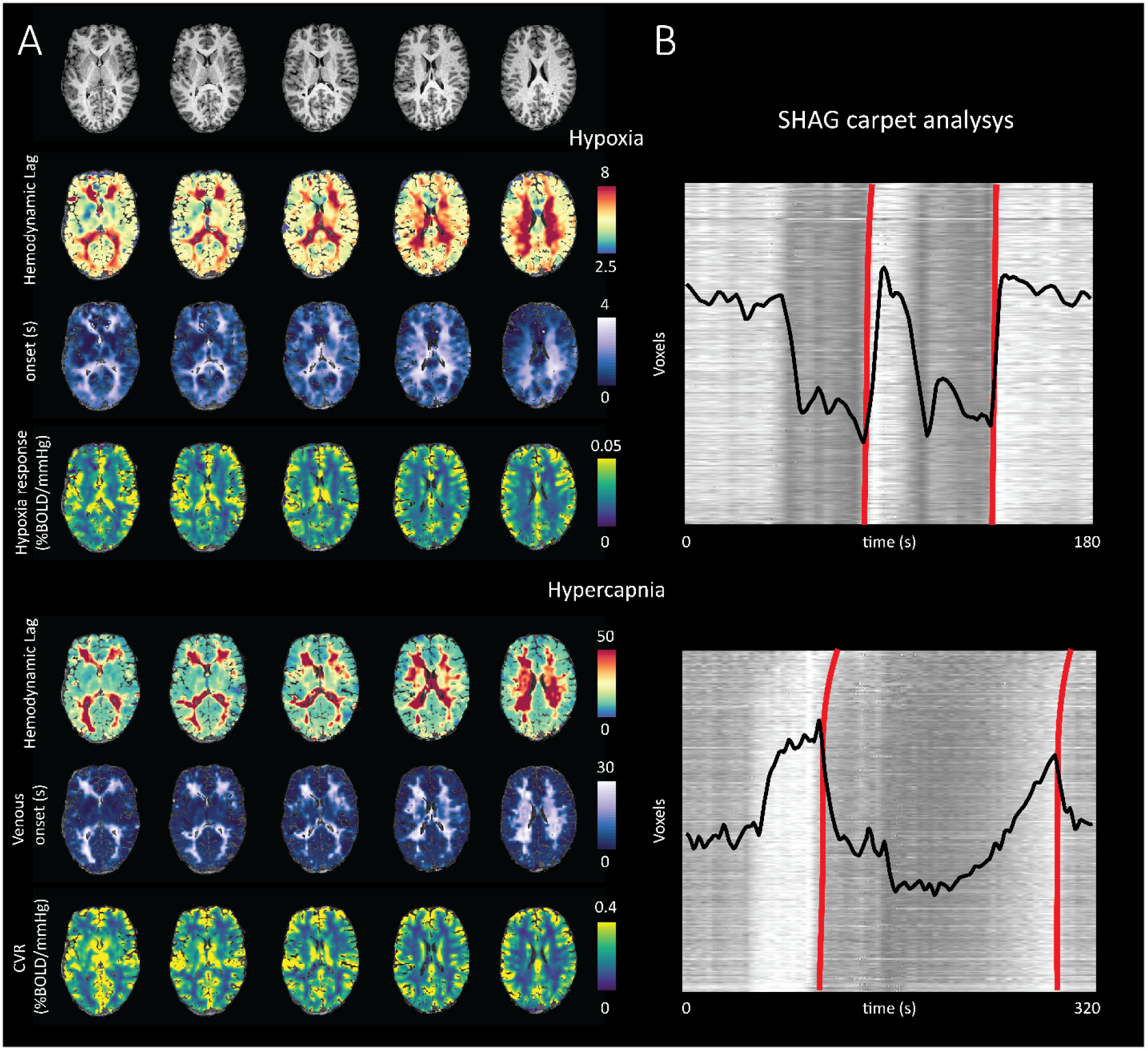
Comparison of hemodynamic parameters generated using hypercapnic versus hypoxic stimulus in a single subject. (A): in general, lag and onset values were considerably longer when based on the hypercapnic stimulus. This is likely due to insensitivity towards arterial signals and true response onset times. The CVR based lag analysis is more highly influenced by vascular resistance mediated flow redistribution, differential sensitivity to CO_2_^45^ and vascular response rate^30^, as well as venous draining topology^37^; (B) carpet plot results fitting the rising edge of the return to normoxia (top) during hypoxic experiments and the falling edge to baseline for hypercapnic experiments (bottom); the descending edge was chosen to do the limited onset contrast generated during the progressive CO_2_ increase. Black traces represent the whole brain BOLD signal responses. Relative to the hypoxic experiment, the hypercapnic response is characterized by greater signal dispersion which translates to longer lag values. Immeasurable CO_2_-modulated arterial properties (i.e. response speed, CO_2_ sensitivity, arterial flow redistribution) translate into longer venous onset times.

### 3.3 hypoxia versus hypercapnia in unilateral carotid artery occlusion

Consistent with the healthy subject example in figure 4, hemodynamic lag for both hypercapnic and hypoxic protocols showed consistent spatial distributions. With hypercapnia as the stimulus, hemodynamic lags were elevated in the healthy hemisphere, suggesting possible inter-hemispheric steal effects. In contrast, the contralateral hemisphere in the hypoxic case showed normal appearing hemodynamic lag (as did most of the GM tissue in the affected hemisphere); this translated to normal onset times (between 0 – 4 s) in the unaffected side but longer onset times in the affected hemisphere. The mean transit time across the brain was calculated at 5 seconds (figure S3), with highest onset values located in the affected hemisphere.

Response characteristics diverged when examining the CVR and hypoxic signal response. As previously determined, hypercapnic stimulus can evoke a vascular steal response, showing widespread re-distribution of blood flow and negative BOLD responses^29^. These pathological responses did manifest in the patient example and compromised the utility of the carpet analysis. The hypoxic signal response remained consistent across hemispheres.

## 4.0 DISCUSSION

In this work, hypoxic respiratory challenges were applied in combination with dynamic BOLD imaging to provide a novel means through which to increase sensitivity weighting towards blood arrival and total blood volume, without the confounding effects related to dynamic vascular responses^30^. Our main finding is that the onset values derived via our modified carpet plot analysis show remarkably consistent temporal distributions that are in line those observed using modalities such as ASL or DSC. This onset, can therefore be considered representative of relative blood arrival time. We report, for the first time, average regional hypoxic response values (expressed as %BOLD/mmHg P_ET_CO_2_) that are representative of relative total blood volume. Finally, comparison with BOLD-CVR responses, dOHb-BOLD signals appear less confounded by dynamic response effects.

The signal onset times reported in our work support the notion that our dOHb-BOLD technique provides a possible alternative means to investigate blood arrival and collateralization of blood flow. Similar to the DSC technique, preliminary work combining dOHb-BOLD have shown promise in delivering similar hemodynamic metrics as their more invasive counterpart. For instance, Vu et al., reported MTT values on the order of 8-9 seconds based on tracer kinetic modeling using a deoxygenated contrast bolus^16^. Using similar methodology, Poublanc et al., reported MTT more in line with physiological ranges expected based on DSC work, at of 3.9 ± 0.6 s and 5.5 ± 0.6 for GM and WM, respectively^31^. Using a hyperoxic bolus, Macdonald et al. were able to derive MTTs of 2.94 ± 0.52 and 3.73 ± 0.60 s for GM and WM despite their insensitivity to arterial contributions. The differences in transit time measured using the hypoxic stimuli may be attributed to differences in the way the hypoxic bolus was administered. Specifically, Poublanc et al. was able to precisely target arterial O_2_ pressure using a computer controlled system.

Where DSC (and dOHb-BOLD) is sensitive to both inflow and outflow of the contrast bolus, ASL-based studies have reported temporal metrics focused mainly on arterial transit times or bolus arrival times. In our method, onset times are reported relative to an internal reference consisting of the first voxels showing the earliest signal onset (i.e. the lowest lines of the carpet plot), AAT and BAT are calculated relative to the time of the labeling pulse. Using multi-PLD pseudo-continuous ASL, Tsujikawa at al., reported increased AAT in affected brain hemispheres of patients with chronic occlusive disease (15 patients: 1.51 ± 0.41 seconds versus and 1.12 ± 0.30 contra-laterally)^32^. Using pulsed ASL, MacIntosh et al., observed significantly longer arrival times in the occipital versus temporal lobes at 0.61 ± 0.096 and 0.935 ± 0.108 seconds, respectively. Interestingly, similar regional differences are seen in our work table 1, along with a clear increases in the affected hemisphere of the patient example (figure 5). Overall, the dOHb-BOLD onset times fall between those reported using ASL (arterial only) and DSC (total vasculature). A strength of dOHb-BOLD over ASL is its sensitivity to WM responses.

**Figure 5:**
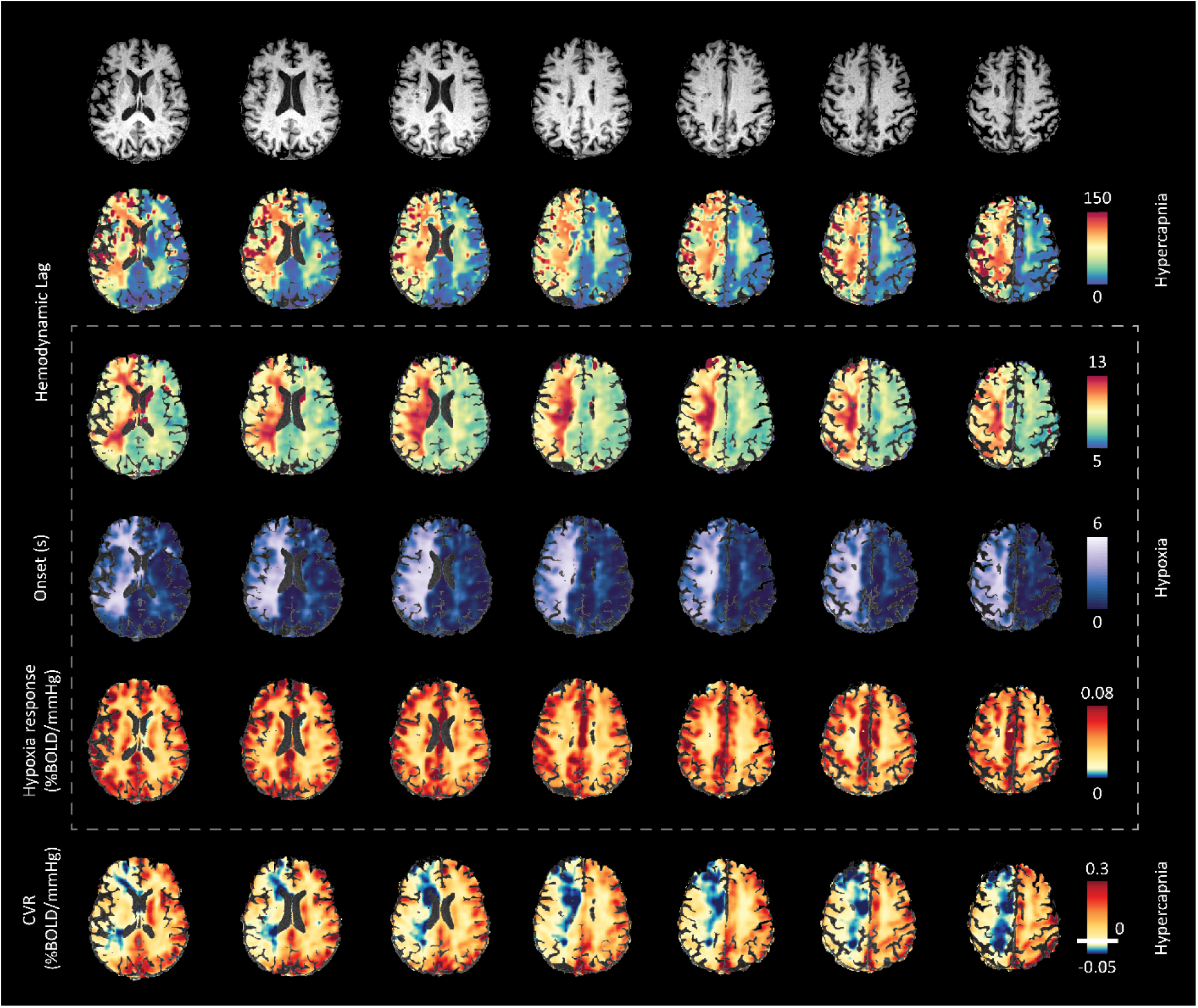
Comparison of hemodynamic parameters generated using hypercapnic versus hypoxic stimulus in a patient with unilateral carotid artery occlusion. Similar to the healthy control case in figure 4, hemodynamic lag distribution was consistent between hypercapnic and hypoxic paradigms (rows 2 and 3). However, CO_2_-mediated lag values where considerably longer and indicated altered flow dynamics in contralateral WM. Unilateral disease can result in bilateral impairment via redistribution effects through collateral networks^46,47^. Onset times derived using the hypoxic protocol remained consistent with controls in the unaffected hemisphere (0 to 4 s; figure 2) while onset time increased in the affected hemisphere (up to 5 s) consistent with presumably elongated transit times associated with collateralization or slower blood flow velocities (carpet plot is provided in figure S2B). Hypoxic response maps (row 5) showed symmetric responses indicating robust blood arrival, while CVR maps (row 6) indicated negative responses related to vascular steal and exhausted vascular reserve capacity in the affected hemisphere. Since these effects generate variable signal responses, impaired reactivity can render the carpet plot analysis unfeasible when using hypercapnia (see figure S2B). *NOTE, CVR color-scale has been adapted compared to figures 3 and 4 in order to highlight negative CVR responses (vascular steal effect) in this patient example.

Dynamic BOLD imaging performed in either the resting-state or throughout hypercapnic gas challenges have also been proposed as methods through which to derive temporal flow information though the assessment of hemodynamic lag^33,34^. A caveat of these approaches is that they derive from vascular responses; either low frequency oscillations of CO_2_-mediated vasodilation (i.e. figure 4/5). The hypercapnia induced response can be slow and depends on the remaining reserve capacity, CO_2_ sensitivity^35^ and vascular response speed^30^. This can drive or long hemodynamic lag in WM regions^18,36^ or the vascular steal phenomenon^29^. While this is may be less apparent when using the resting state signal, a hindrance to robust lag estimation arises due to the low CNR or blood-borne signals. Moreover, signals are subject to additional confounding contributions related to cerebral draining architecture which can influence local dHb concentrations^37^ and obscure true blood arrival or transit times. Finally, resting-state and hypercapnia/hyperoxic BOLD signal changes are localized in veins. This means that these techniques are insensitive to almost the entirety of the arterial vascular network, and so can never provide accurate information on blood arrival time, transit time or collateral blood flow as what are provided using ASL, DCE and now potentially dOHb-BOLD.

A further finding of interest is that dOHb-BOLD can provide a surrogate measure of total blood volume. Vascular volume fraction as measured using ^15^O PET has been reported as approximately 1.93 fold higher in GM (5.2 ± 1.4%) versus WM (2.7 ± 0.6%) which is similar with our findings of a 1.7 fold increase in hypoxic signal response between GM and deep WM^28^. A potential confound to this application is that prolonged hypoxia is known to have a vasoactive effect leading to increased CBF^38,39^ (and CVR). However, the response dynamics seem more sluggish when compared to commonly used vasodilators such as carbon dioxide. The work of Harris et al.^40^, reported a 15.4% increase in hypoxia-modulated CBF, albeit with an CBF onset delay of 185 s when transitioning from normoxia to hypoxia and a non-significant delay when returning to baseline arterial oxygen pressure. This considerable delay reinforces the assumption that the CBF response, when transitioning towards hypoxia in our study, is unlikely to lead to significant hemodynamic effects and supports our claims of measuring true response onset (or relative blood arrival times) and potentially, total blood volume. In line with the results of Harris et al., we also observed more rapid transitions towards normoxia than towards hypoxia (figure S2), which guided our decision to quantify onset using rising, rather than falling edges during the carpet plot analysis. This asymmetric response may be explained by the presence of more plasma-dissolved O_2_ during the baseline state that was available to bind to hemoglobin during the initial transition to hypoxia. The temporal asymmetry may also arise due to the fact that RespirAct system requires multiple breathing cycles to establish hypoxia, but is able to transition back to normoxia within one breath.

## 5.0 CONSIDERATIONS

The main consideration of our work relates to the use of hemodynamic lag values to sort carpet plot voxels, meaning that the onset time may be weighted by the accuracy of the lag calculation. This could lead to errors in cases where voxels are temporally mislabeled (i.e. improperly ordered with respect to the edge calculation). Our preliminary comparison suggests that this approach is more reliable for hypoxia as compared to hypercapnia do the obvious sharper transitions and lack of dynamic response effects when transitioning between physiological states (see figure S3). Alternative ways of sorting (particularly for CO_2_-mediated changes) could be based on signal dispersion (defined as ‘Tau’ in the work of Poublanc et al.^30^). Using this approach, onset times could then also be fit explicitly using a gamma variate function^37^. Nevertheless, regions with limited (i.e. deep WM) or impaired CVR will not provide any meaningful information with respect to arrival time when using a CO_2_ stimulus. In pathological cases exhibiting vascular steal phenomenon, the inverse polarity of the signal response in steal versus healthy regions will hinder the edge detection due to a mixing of rising and falling responses (see figure S3B).

In the original work of Fitzgerald et al., reported transit times when using BOLD data were representative of venous transit time (assumed as whole brain). In our updated method using dOHb-BOLD, we are mainly sensitive to arterial onset (i.e. entry of the bolus into arterial voxels) since we cannot distinguish between arterial and venous signals. It is expected that the dOHb bolus would induce a bi-phasic response behavior; an initial peak as de-oxygenated (SaO_2_ approx. 75%) blood enters arteries, followed by a second peak as the hypoxic bolus enters the venous system. Based on MRI measurements of resting venous saturation (S_v_O_2_) using T2-Relaxation-Under-Spin-Tagging (TRUST)^41^ MRI, Deckers et al.^42^ reported S_v_O_2_ on the order of 60-70% while breathing room air. Under the assumption that hypoxic breathing is iso-metabolic, S_V_O_2_ under hypoxia could be expected to drop to as little as 35-45%. This could manifest in a significant BOLD signal differences emanating from veins versus arteries. Exploring this hypothesis would require a considerably higher sampling rate and higher spatial resolution than what was used in the current study with an additional S_V_O_2_ measurement to facilitate more in-depth physiological modelling. To investigate this, future experimental designed might include the use of ultra-high field MRI in combination with surface receive coils^43^ or fast line-scanning approaches^44^. Of further interest could be an in-depth of hypoxic arrival time maps with perfusion metrics derived from the ‘gold-standard’ DSC method. Additionally, a systematic comparison between temporal characteristics derived via hypercapnic versus hypoxic (and perhaps hyperoxic) experiments is warranted to confirm our interpretations made based on data presented in figures 4 and 5. From a clinical perspective, the combination of our modified carpet plot analysis with dOHb-BOLD can provide insight into the status of collateral flow networks. As shown in our preliminary patient example, dOHb-BOLD onset times reflect normal baseline perfusion even in the presence of vascular steal. The addition of dOHb-BOLD to clinical CVR studies may help to better stratify the severity of vascular impairment due to disease. This warrants a deeper examination of our method applied in patients with different cerebrovascular pathology.

## Conclusion

Our dOHb-BOLD method can be used to quantify the onset of immediate signal changes to provide a measure of arterial blood arrival time. In cases with exhausted reserve capacity or confounding flow effects such as vascular steal, dOHb-BOLD can potentially inform collateral flow pathways providing a valuable compliment to clinical CVR measures.

## Declarations of interest

JAF, DJM contributed to the development of the automated end-tidal targeting device, RespirAct™ (Thornhill Research Inc., TRI) used in this study and have equity in the company. RespirAct™ is a non-commercial device built by TMI to enable measurement of CVR in scientific studies. OS and JD receive salary support from TRI. TRI provided no other support for the study. All other authors have no disclosures to report.

## Data and code availability

The analysis tools used to generate the results presented in this manuscript are freely available via the open-source seeVR toolbox (https://github.com/abhogal-lab/seeVR). MRI and physiological data can be made available based on the submission of a formal project outline. Anonymized data will be shared by request from any qualified investigator for purposes such as replicating procedures and results presented in the article, provided that data transfer is in agreement with the University Health Network and Health Canada legislation on the general data protection regulation

## Funding

This work was supported by a Dutch research council talent grant awarded to Alex A. Bhogal (NWO VENI: *The ischemic fingerprint*, [VI.VENI.194.056].

## REFERENCES

1. van Osch MJ, Teeuwisse WM, Chen Z, Suzuki Y, Helle M, Schmid S. Advances in arterial spin labelling MRI methods for measuring perfusion and collateral flow. Journal of cerebral blood flow and metabolism : official journal of the International Society of Cerebral Blood Flow and Metabolism. 2018;38(9):1461–1480.

2. McVerry F, Liebeskind DS, Muir KW. Systematic Review of Methods for Assessing Leptomeningeal Collateral Flow. American Journal of Neuroradiology. 2012;33(3):576–582.

3. Hartkamp NS, Bokkers RP, van der Worp HB, van Osch MJ, Kappelle LJ, Hendrikse J. Distribution of cerebral blood flow in the caudate nucleus, lentiform nucleus and thalamus in patients with carotid artery stenosis. European radiology. 2011;21(4):875–881.

4. Sebök M, Niftrik C, Lohaus N, et al. Leptomeningeal collateral activation indicates severely impaired cerebrovascular reserve capacity in patients with symptomatic unilateral carotid artery occlusion. Journal of cerebral blood flow and metabolism : official journal of the International Society of Cerebral Blood Flow and Metabolism. 2021;41(11):3039–3051.

5. Sebök M, Esposito G, Niftrik C, et al. Flow augmentation STA-MCA bypass evaluation for patients with acute stroke and unilateral large vessel occlusion: a proposal for an urgent bypass flowchart. Journal of neurosurgery. 2022:1–9.

6. Okell TW, Harston GWJ, Chappell MA, Sheerin F, Kennedy J, Jezzard P. Measurement of collateral perfusion in acute stroke: a vessel-encoded arterial spin labeling study. Scientific reports. 2019;9(1):8181.

7. Zaharchuk G. Arterial Transit Awesomeness. Radiology. 2020;297(3):661–662.

8. Lou X, Yu S, Scalzo F, et al. Multi-delay ASL can identify leptomeningeal collateral perfusion in endovascular therapy of ischemic stroke. Oncotarget. 2017;8(2):2437–2443.

9. Paling D, Thade Petersen E, Tozer DJ, et al. Cerebral arterial bolus arrival time is prolonged in multiple sclerosis and associated with disability. Journal of cerebral blood flow and metabolism : official journal of the International Society of Cerebral Blood Flow and Metabolism. 2014;34(1):34–42.

10. Buxton RB, Frank LR, Wong EC, Siewert B, Warach S, Edelman RR. A general kinetic model for quantitative perfusion imaging with arterial spin labeling. Magnetic resonance in medicine. 1998;40(3):383–396.

11. Seiler A, Brandhofe A, Gracien RM, et al. DSC perfusion-based collateral imaging and quantitative T2 mapping to assess regional recruitment of leptomeningeal collaterals and microstructural cortical tissue damage in unilateral steno-occlusive vasculopathy. Journal of cerebral blood flow and metabolism : official journal of the International Society of Cerebral Blood Flow and Metabolism. 2021;41(1):67–81.

12. Sobczyk O, Sam K, Mandell DM, et al. Cerebrovascular Reactivity Assays Collateral Function in Carotid Stenosis. Frontiers in physiology. 2020;11:1031.

13. Duffin J, Sobczyk O, Crawley A, et al. The role of vascular resistance in BOLD responses to progressive hypercapnia. Human Brain Mapping. 2017;38(11):5590–5602.

14. Donahue MJ, Faraco CC, Strother MK, et al. Bolus arrival time and cerebral blood flow responses to hypercarbia. Journal of cerebral blood flow and metabolism : official journal of the International Society of Cerebral Blood Flow and Metabolism. 2014;34(7):1243–1252.

15. Poublanc J, Sobczyk O, Shafi R, et al. Perfusion MRI using endogenous deoxyhemoglobin as a contrast agent: Preliminary data. Magnetic resonance in medicine. 2021;86(6):3012–3021.

16. Vu C, Chai Y, Coloigner J, et al. Quantitative perfusion mapping with induced transient hypoxia using BOLD MRI. Magnetic resonance in medicine. 2021;85(1):168–181.

17. Fitzgerald B, Yao JF, Talavage TM, Hocke LM, Frederick BD, Tong Y. Using carpet plots to analyze transit times of low frequency oscillations in resting state fMRI. Scientific reports. 2021;11(1):7011.

18. Champagne AA, Bhogal AA. Insights Into Cerebral Tissue-Specific Response to Respiratory Challenges at 7T: Evidence for Combined Blood Flow and CO(2)-Mediated Effects. Frontiers in physiology. 2021;12:601369.

19. Frederick B, Nickerson LD, Tong Y. Physiological denoising of BOLD fMRI data using Regressor Interpolation at Progressive Time Delays (RIPTiDe) processing of concurrent fMRI and near-infrared spectroscopy (NIRS). Neuroimage. 2012;60(3):1913–1923.

20. Donahue MJ, Strother MK, Lindsey KP, Hocke LM, Tong Y, Frederick BD. Time delay processing of hypercapnic fMRI allows quantitative parameterization of cerebrovascular reactivity and blood flow delays. Journal of cerebral blood flow and metabolism : official journal of the International Society of Cerebral Blood Flow and Metabolism. 2016;36(10):1767–1779.

21. Bhogal AA. seeVR: a toolbox for analyzing cerebrovascular reactivity data. Zenodo. 2021;(v1.01). 22.

22. Jenkinson M, Beckmann CF, Behrens TE, Woolrich MW, Smith SM. FSL. Neuroimage. 2012;62(2):782–790.

23. Jenkinson M, Smith S. A global optimisation method for robust affine registration of brain images. Med Image Anal. 2001;5(2):143–156.

24. Smith SM. Fast robust automated brain extraction. Hum Brain Mapp. 2002;17(3):143–155.

25. Zhang Y, Brady M, Smith S. Segmentation of brain MR images through a hidden Markov random field model and the expectation-maximization algorithm. IEEE transactions on medical imaging. 2001;20(1):45–57.

26. Severinghaus JW. Simple, accurate equations for human blood O2 dissociation computations. Journal of Applied Physiology. 1979;46(3):599–602.

27. Balaban DY, Duffin J, Preiss D, et al. The in-vivo oxyhaemoglobin dissociation curve at sea level and high altitude. Respiratory Physiology & Neurobiology. 2013;186(1):45–52.

28. Leenders KL, Perani D, Lammertsma AA, et al. Cerebral blood flow, blood volume and oxygen utilization. Normal values and effect of age. Brain. 1990;113 (Pt 1):27–47.

29. Sobczyk O, Battisti-Charbonney A, Fierstra J, et al. A conceptual model for CO2-induced redistribution of cerebral blood flow with experimental confirmation using BOLD MRI. Neuroimage. 2014;92:56–68.

30. Poublanc J, Crawley AP, Sobczyk O, et al. Measuring cerebrovascular reactivity: the dynamic response to a step hypercapnic stimulus. Journal of cerebral blood flow and metabolism : official journal of the International Society of Cerebral Blood Flow and Metabolism. 2015;35(11):1746– 1756.

31. Ibaraki M, Ito H, Shimosegawa E, et al. Cerebral Vascular Mean Transit Time in Healthy Humans: A Comparative Study with PET and Dynamic Susceptibility Contrast-Enhanced MRI. Journal of Cerebral Blood Flow & Metabolism. 2006;27(2):404–413.

32. Tsujikawa T, Kimura H, Matsuda T, et al. Arterial Transit Time Mapping Obtained by Pulsed Continuous 3D ASL Imaging with Multiple Post-Label Delay Acquisitions: Comparative Study with PET-CBF in Patients with Chronic Occlusive Cerebrovascular Disease. PLoS One. 2016;11(6):e0156005–e0156005.

33. Aso T, Jiang G, Urayama S-i, Fukuyama H. A Resilient, Non-neuronal Source of the Spatiotemporal Lag Structure Detected by BOLD Signal-Based Blood Flow Tracking. Front Neurosci. 2017;11.

34. Tong Y, Frederick Bd. Tracking cerebral blood flow in BOLD fMRI using recursively generated regressors. Human Brain Mapping. 2014;35(11):5471–5485.

35. Bhogal AA, Siero JC, Fisher JA, et al. Investigating the non-linearity of the BOLD cerebrovascular reactivity response to targeted hypo/hypercapnia at 7T. Neuroimage. 2014;98:296–305.

36. Bhogal AA, Philippens ME, Siero JC, et al. Examining the regional and cerebral depth-dependent BOLD cerebrovascular reactivity response at 7T. Neuroimage. 2015;114:239–248.

37. Bhogal AA. Medullary vein architecture modulates the white matter BOLD cerebrovascular reactivity signal response to CO2: Observations from high-resolution T2* weighted imaging at 7T. Neuroimage. 2021;245:118771.

38. Jensen MLF, Vestergaard MB, Tønnesen P, Larsson HBW, Jennum PJ. Cerebral blood flow, oxygen metabolism, and lactate during hypoxia in patients with obstructive sleep apnea. Sleep. 2018;41(3).

39. Cohen PJ, Alexander SC, Smith TC, Reivich M, Wollman H. Effects of hypoxia and normocarbia on cerebral blood flow and metabolism in conscious man. Journal of Applied Physiology. 1967;23(2):183–189.

40. Harris AD, Murphy K, Diaz CM, et al. Cerebral blood flow response to acute hypoxic hypoxia. NMR in biomedicine. 2013;26(12):1844–1852.

41. Lu H, Ge Y. Quantitative evaluation of oxygenation in venous vessels using T2-Relaxation-Under-Spin-Tagging MRI. Magnetic resonance in medicine. 2008;60(2):357–363.

42. Deckers PT, Bhogal AA, Dijsselhof MB, et al. Hemodynamic and metabolic changes during hypercapnia with normoxia and hyperoxia using pCASL and TRUST MRI in healthy adults. Journal of Cerebral Blood Flow & Metabolism.0(0):0271678X211064572.

43. Schellekens W, Bhogal AA, Roefs ECA, Báez-Yáñez MG, Siero JCW, Petridou N. The many layers of BOLD. On the contribution of different vascular compartments to laminar fMRI. bioRxiv. 2021:2021.2010.2021.465359.

44. Raimondo L, Knapen T, Oliveira ĺAF, et al. A line through the brain: implementation of human line-scanning at 7T for ultra-high spatiotemporal resolution fMRI. Journal of Cerebral Blood Flow & Metabolism. 2021;41(11):2831–2843.

45. Thomas BP, Liu P, Park DC, van Osch MJP, Lu H. Cerebrovascular Reactivity in the Brain White Matter: Magnitude, Temporal Characteristics, and Age Effects. Journal of Cerebral Blood Flow & Metabolism. 2013;34(2):242–247.

46. Deckers PT, van Hoek W, Kronenburg A, et al. Contralateral improvement of cerebrovascular reactivity and TIA frequency after unilateral revascularization surgery in moyamoya vasculopathy. NeuroImage: Clinical. 2021;30:102684.

47. Sam K, Poublanc J, Sobczyk O, et al. Assessing the effect of unilateral cerebral revascularisation on the vascular reactivity of the non-intervened hemisphere: a retrospective observational study. BMJ Open. 2015;5(2):e006014.

