## Supplementary material for "Quantifying cerebral blood arrival times using hypoxia-mediated arterial BOLD contrast": figure S

**SUPLEMENTARY FIGURES**

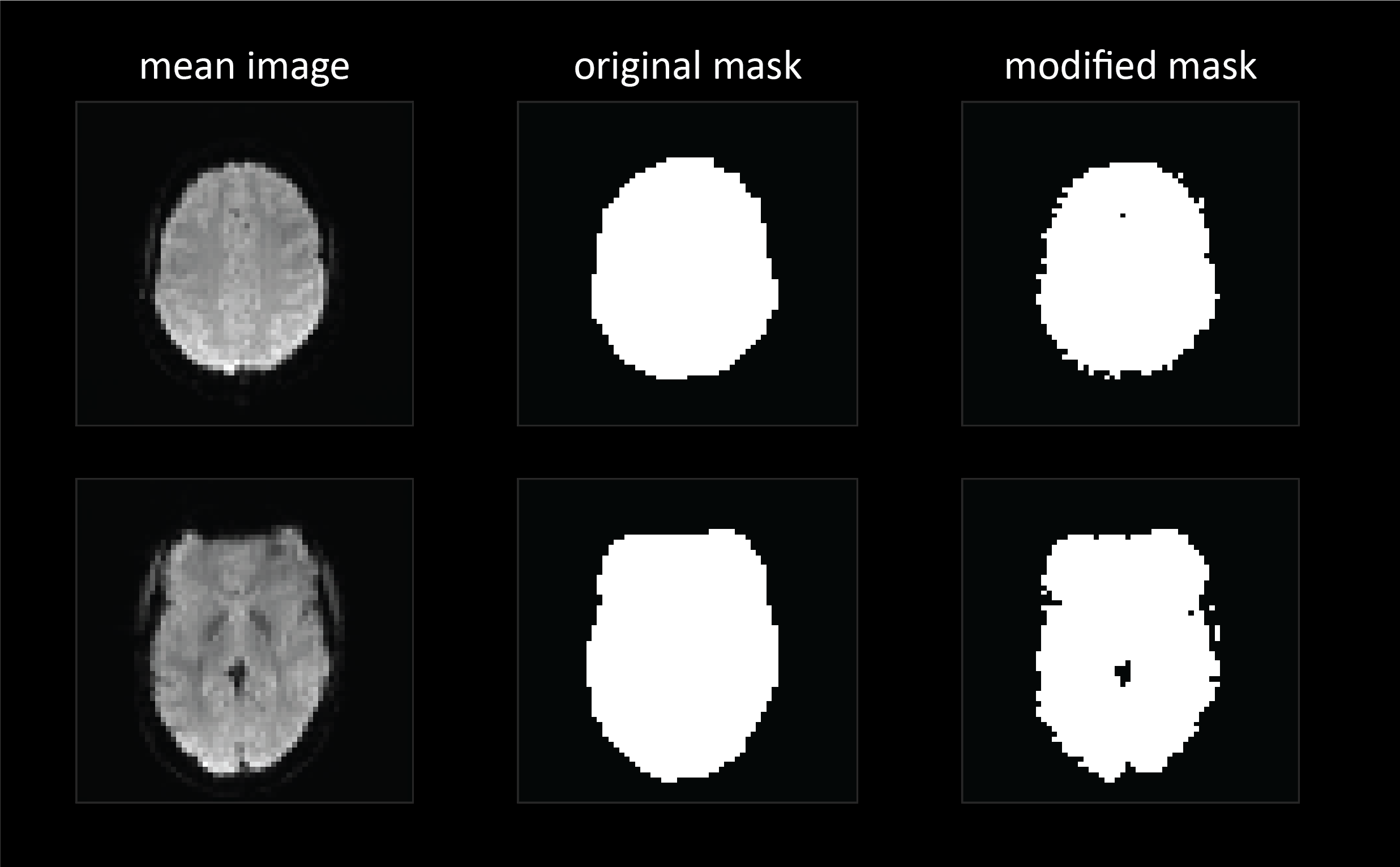

Figure S1: ***Removal of signal contributions from large vessels.*** Large veins cause extravascular signal effects that can translate high CVR values that may not be reflective of the true vascular response and may obscure arterial information. An refined mask is created to ignore large vessel and CSF voxels.

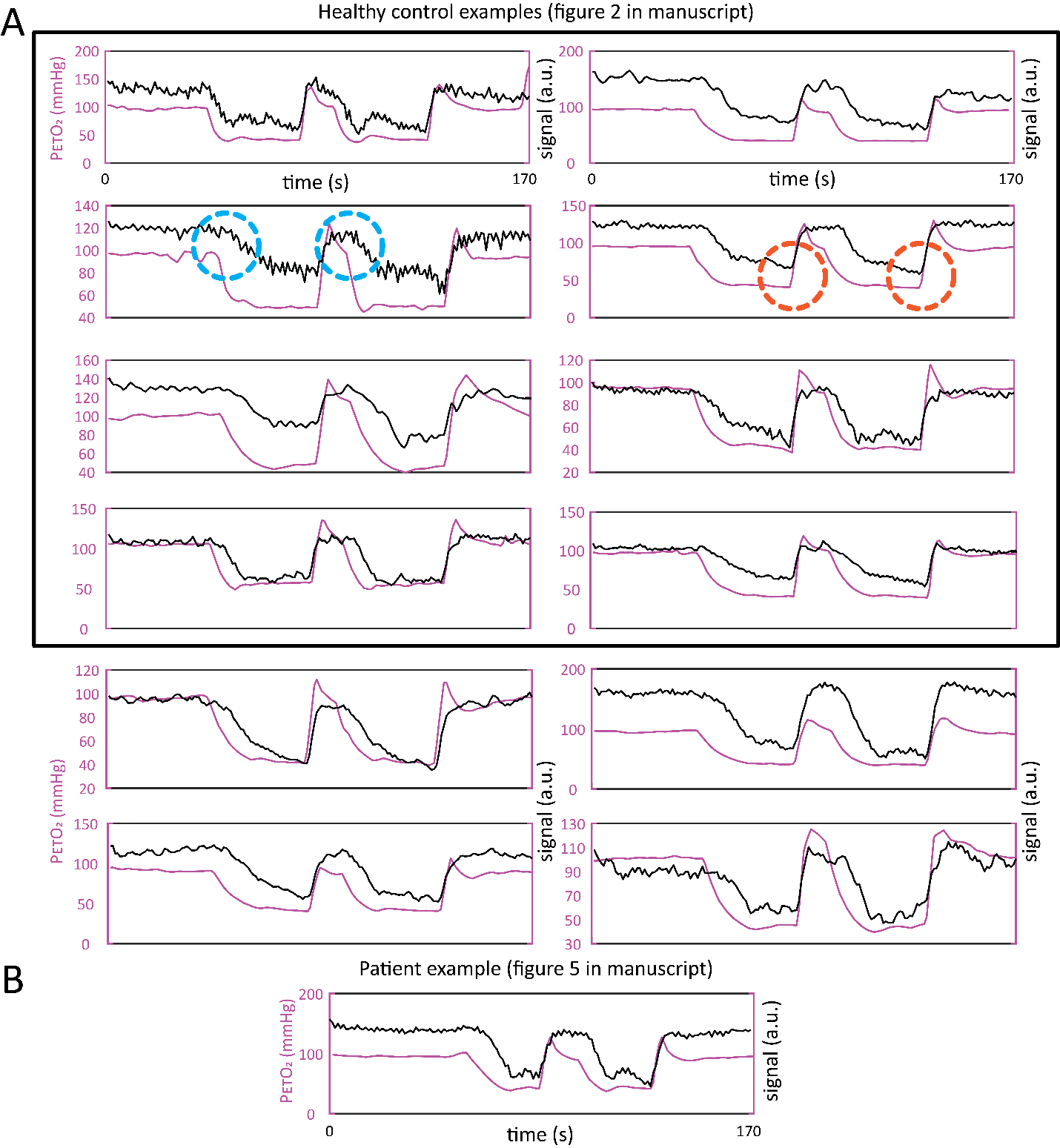

Figure S2: ***Bulk-alignment of raw average GM BOLD signal trace (in arbitrary MR scanner units) with resampled (to TR) P_ET_O_2_ traces for all 12 healthy controls (A) and the single CAO patient (B).*** Data contained within the black bounding box is derived from the subjects shown in figure 2 of the main manuscript. In general, the hypoxic stimulus targeted at 40 mmHg lead to robust BOLD signal changes in all subjects.Of note is the dOHb-BOLD signal response lag (denoted by blue circles) when going from normoxia to hypoxia as compared to the immediate and sharp transition from going from hypoxia to normoxia (denoted by orange circles).

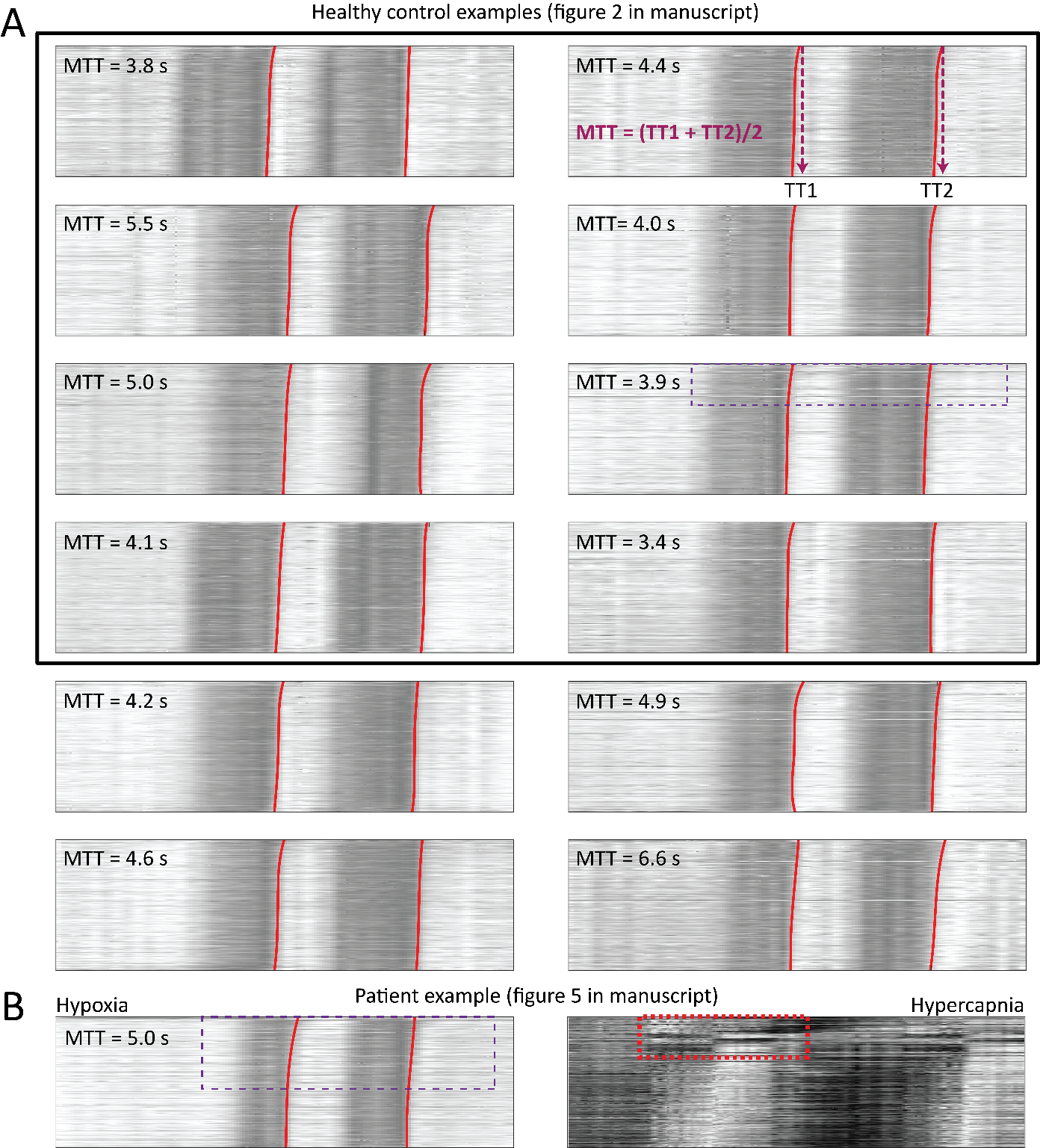

Figure S3: ***Carpet plot analysis with fitted transit/onset edges for all 12 healthy controls (A) and the single CAO patient (B)*.** The MTT calculated by averaging the values derived from both fitted edges is provided for each subject (inset). Healthy subject edges showed mainly linear behavior until only the top (in WM) region of the carpet plot (see hatched box in S3A) while for the CAO patient, the longer onset times in the affected hemisphere lead to increase non-linear behavior (hatched box S3B: left). This lead to an increase in the average MTT as compared with the average control value. The hypercapnic plot (3SB: right) is zoomed to highlight the upper region of the plot. Long delays and vascular steal effects, visible as a signal inversion, prevent accurate fitting of the response edge. Note the longer delays.
